# Biomarker model for cancer: Development of fast LC-MS/MS method for reduced and oxidized glutathione

**DOI:** 10.1101/2020.04.26.058412

**Authors:** Ray F. Nassar, Praneeth Chitralia, Roger Rushworth, Charles O’Donnell, Iswarya Gagrin

**Affiliations:** Engineering and Science University Magnet School, 500 Boston Post Road, West Haven, CT, USA; Chemistry & Chemical Engineering Department University of New Haven, 300 Boston Post Rd, West Haven, CT, USA

## Abstract

In recent years, there have been significant efforts devoted to countering the challenge of detecting cancer in early stages. Reduced glutathione (GSH) plays an important role in the antioxidant system and is required for the maintenance of the redox status of the cell, defense against free radicals and detoxification of toxic compounds. GSH may be converted to oxidized glutathione (GSSG) during the oxidative stress that it regularly undergoes when combating cancer cells. Therefore, the ratio of GSH to the total amount of glutathione can be an extremely useful biomarker for detecting cancer. However, there has yet to be an effective method of detecting and quantifying glutathione in cells, making it extremely difficult to use as a biomarker. In this study, we have created an effective method of detecting both forms of glutathione, utilizing high-performance liquid chromatography-tandem mass spectrometry (HPLC-MS/MS). The analysis time took less than 1 minute, and we were able to quantify both GSH and GSSG in one method. The limit of quantitation is 1 ng/mL, and we ran three trials, each examining a range of concentrations, from 1 to 500 ng/ml of GSH and GSSG. Results were calculated using peak area ratios, using HPLC-MS/MS technology, we were able to determine both the amounts of GSH and GSSG in a single method, creating a fast, reliable, non-invasive, and cost-effective method of testing early stages of cancer.

## Introduction

In normal conditions, cells will have glutathione, an antioxidant, which engages in a variety of functions, most involved in maintaining cellular redox homeostasis. Glutathione is the prevalent antioxidant in mammals, normally having a concentration of 1-10 mM. Reduced glutathione (GSH) also has an inactive, oxidized version (GSSG) (Figure 1). The chemical structures formula of both types of glutathione (GSH, GSSG).

**Figure 1.**
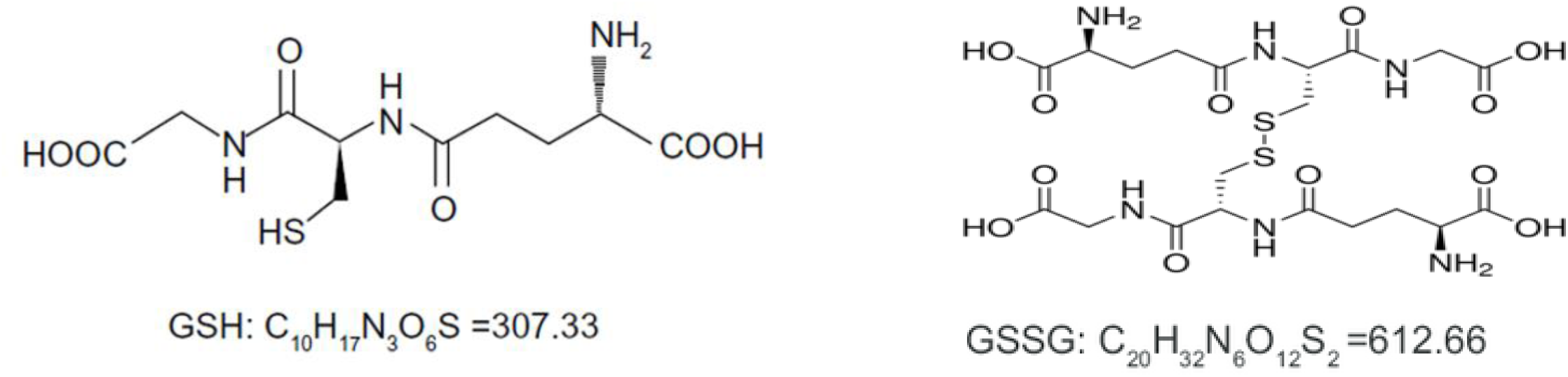
The skeletal formulae and structures of both types of glutathione (GSH, GSSG)

In cancer cells, there is a higher amount of antioxidants in cells than standard. Cells maintain a balance of antioxidants (GSH) against Reactive oxygen species generated by the different physiological processes. (Ankita Bansal and M. Celeste Simon, J. Cell Biol. 2018 Vol. 217 No. 7). During the oxidative stress that cells endure while countering cancer, reduced glutathione (GSH) may be converted to its oxidized version (GSSG) (Figure 2).

**Figure 2.**
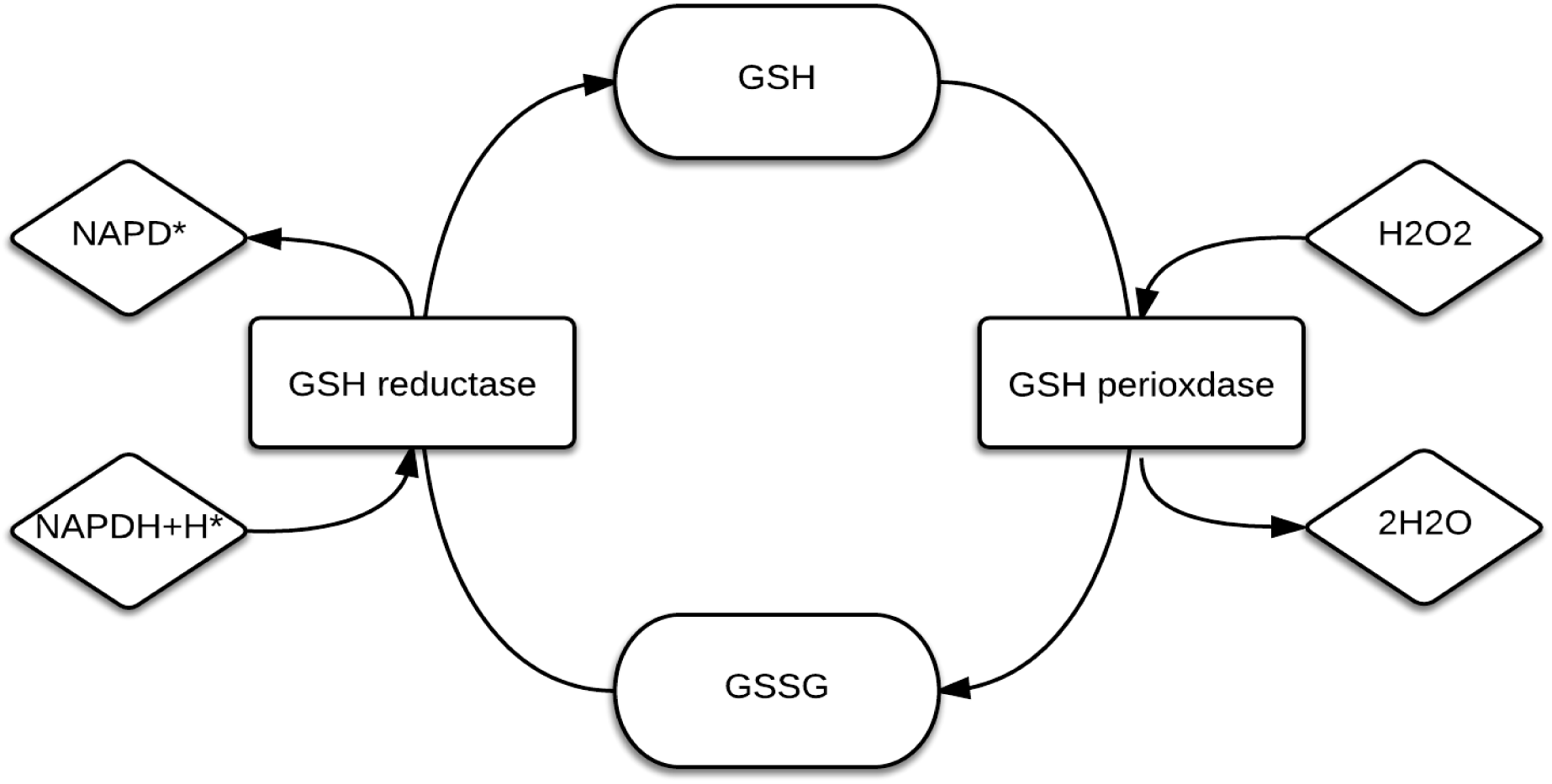
The cycle that GSH and GSSG go through to be converted to one another

GSH and GSSG have been measured by a number of methods. Although several methods are available for measuring GSH and GSSG, all have disadvantages including the need to generate derivatives, the inability to conveniently measure both GSH and GSSG, and lack of enough sensitivity to allow detection in small samples. We have developed a simple and sensitive liquid chromatography–tandem mass spectrometry (LC–MS/MS) method for GSH and GSSG. The assay, a fast and reliable method to measure oxidized and reduced form of glutathione was developed.

### Principles of the method

This method was designed to detect GSH and GSSG using LC-MS/MS. The standard curve range was 5-1000 ng/mL.

### Materials

Water, Ethanol, Formic Acid, GSH, GSSG (all bought by Sigma Aldrich)

### Instrumentation

LCMS (Sciex API 3000 triplequad and LC-Agilent)

### Preparation of Stock Standard Solutions

Stock standard solutions for glutathione reduced form and glutathione oxidized form are prepared in duplicate and compared using the appropriate weight for the lot purity, moisture, and salt correction.

As shown in table 1, place 200 µL of the standard stock solution into a polypropylene vial and add 800 µL of the diluent to the removed standard stock solution amount. The glutathione (reduced and oxidized) concentration in this solution [Std5] will be 1000.00 ng/mL. Place 200 µL of Std5 into a separate vial and add 800 µL of the diluent to Std5, making the concentration of the solution [Std4] 200.00 ng/mL of glutathione. Place 100 µL of Std4 into another separate vial and add 300 µL of diluent to the removed Std4. The solution’s [Std3] concentration should then be 50.00 ng/mL of glutathione. Place 100 µL of Std3 into another separate vial and add 400 µL of diluent to the removed Std3; the glutathione concentration of the solution [Std2] will be 10.00 ng/mL. Place 100 µL of Std2 into another separate vial and add 100 µL of diluent to the removed Std2. The glutathione concentration of the solution [Std1] will be 5.00 ng/mL. Place 25 µL of Std1 into another separate vial and add 100 µL of the diluent to the removed Std1, making the final solution’s concentration 1.00 ng/mL of glutathione. All solutions are stored in polypropylene vials in a refrigerator set to maintain 2°C to 8°C.

**Table 1.**
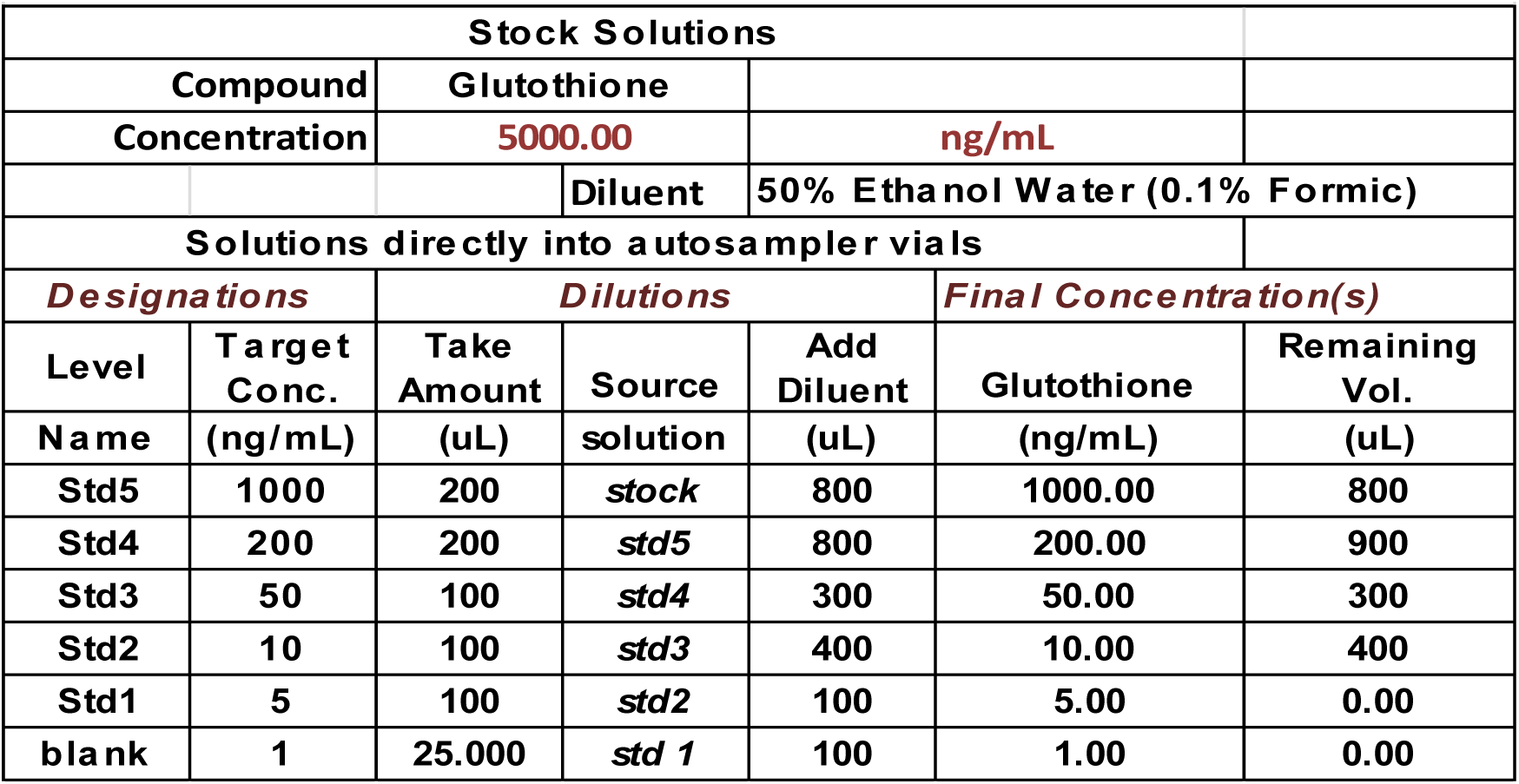
Calibrator Prep Chart

**Figure 3.**
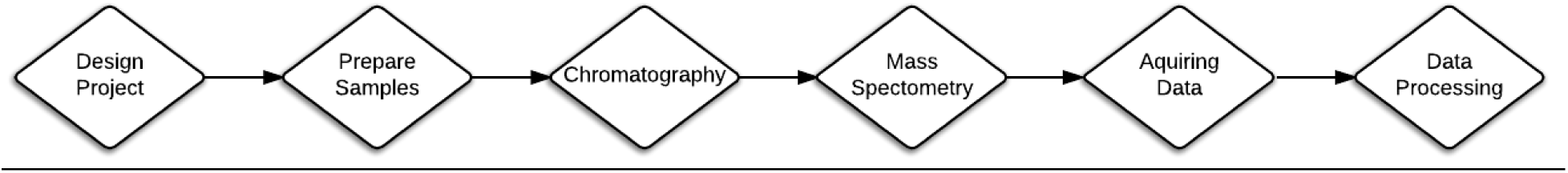
A flowchart depicting numerous steps in the method

## Results

In Figure 4, a representative MS spectrum of GSH is shown, used to determine an ion and its intensity, from glutathione solution. Glutathione has m/z mass to charge of 307. However, this LC-MS method lacks specificity.

**Figure 4.**
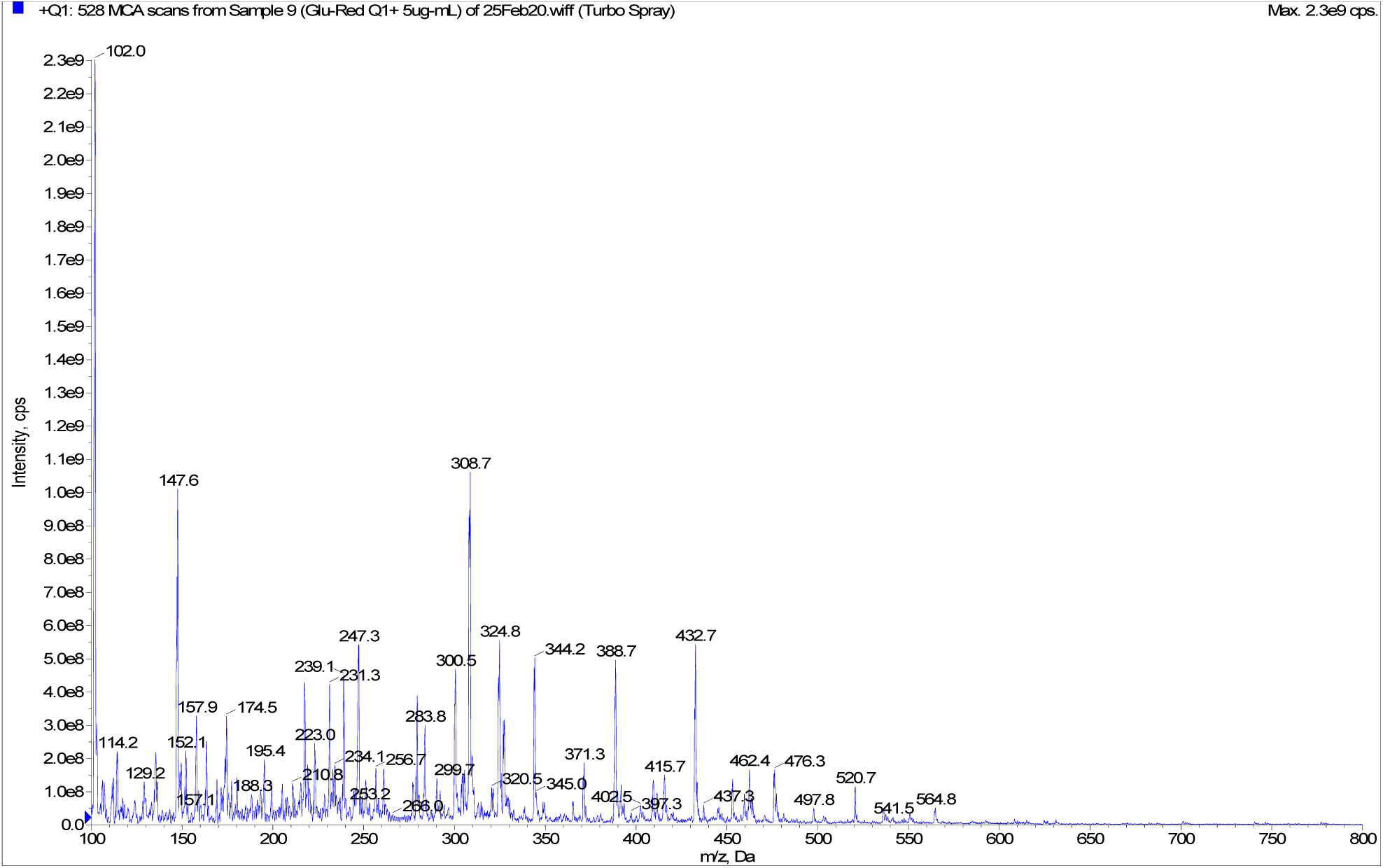
MS spectra for GSH

In Figure 5, a representative MS spectrum of GSSG is shown, used to determine an ion and its intensity, from glutathione solution. Glutathione has m/z mass to charge of 612.

**Figure 5.**
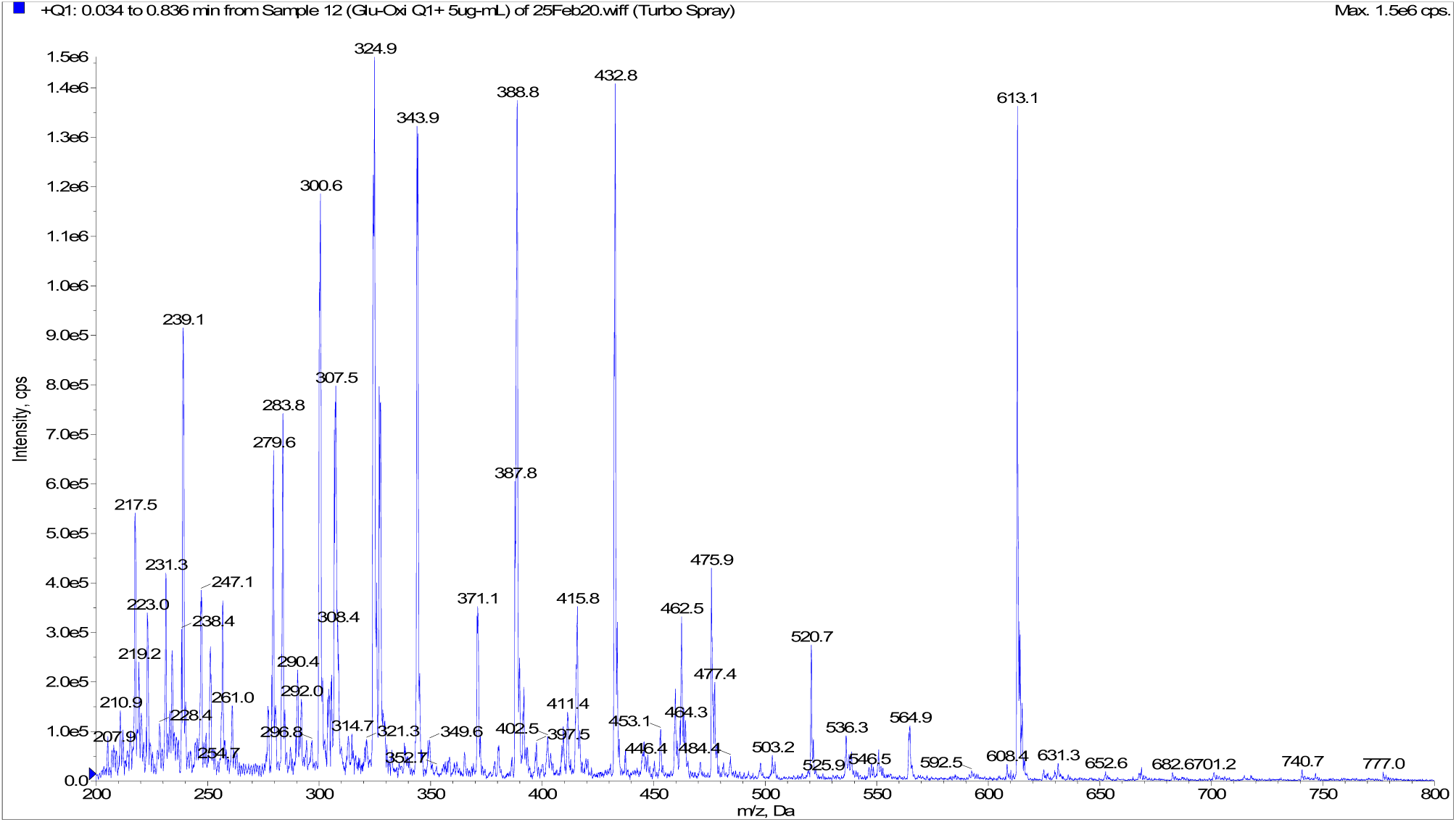
MS spectra for GSSG

### MS/MS methods

A summary of the MS/MS method for GSH and GSSG is shown in table 2. The transition monitored for GSH m/z 307 to 179. The transition monitored for GSSG m/z 612 to 483. The collision energy 20eV for both GSH and GSSG.

**Table 2.** Summary of MS/MS parameters for GSH and GSSG

| Compound ID | Compound Name | Transition Monitored | Dwell Time (msec) | Collision Energy (eV) | Approximate Retention Time (min) |
| --- | --- | --- | --- | --- | --- |
| Analyte 1 | GSH | 307→179 | 100 | 20 | 0.2 |
| Analyte 2 | GSSG | 612→483 | 100 | 20 | 0.2 |

In Table 2, the MS/MS method that we have developed for determining GSH and GSSG is shown. We determined from the MS spectrums that GSH and GSSG had 307 and 612 ions respectively, and we used the LCMS/MS method to break these ions, and test for the amount of product ions, giving us 179 product ions of GSH, and 483 for GSSG (Figures 6, 7).

**Figure 6.**
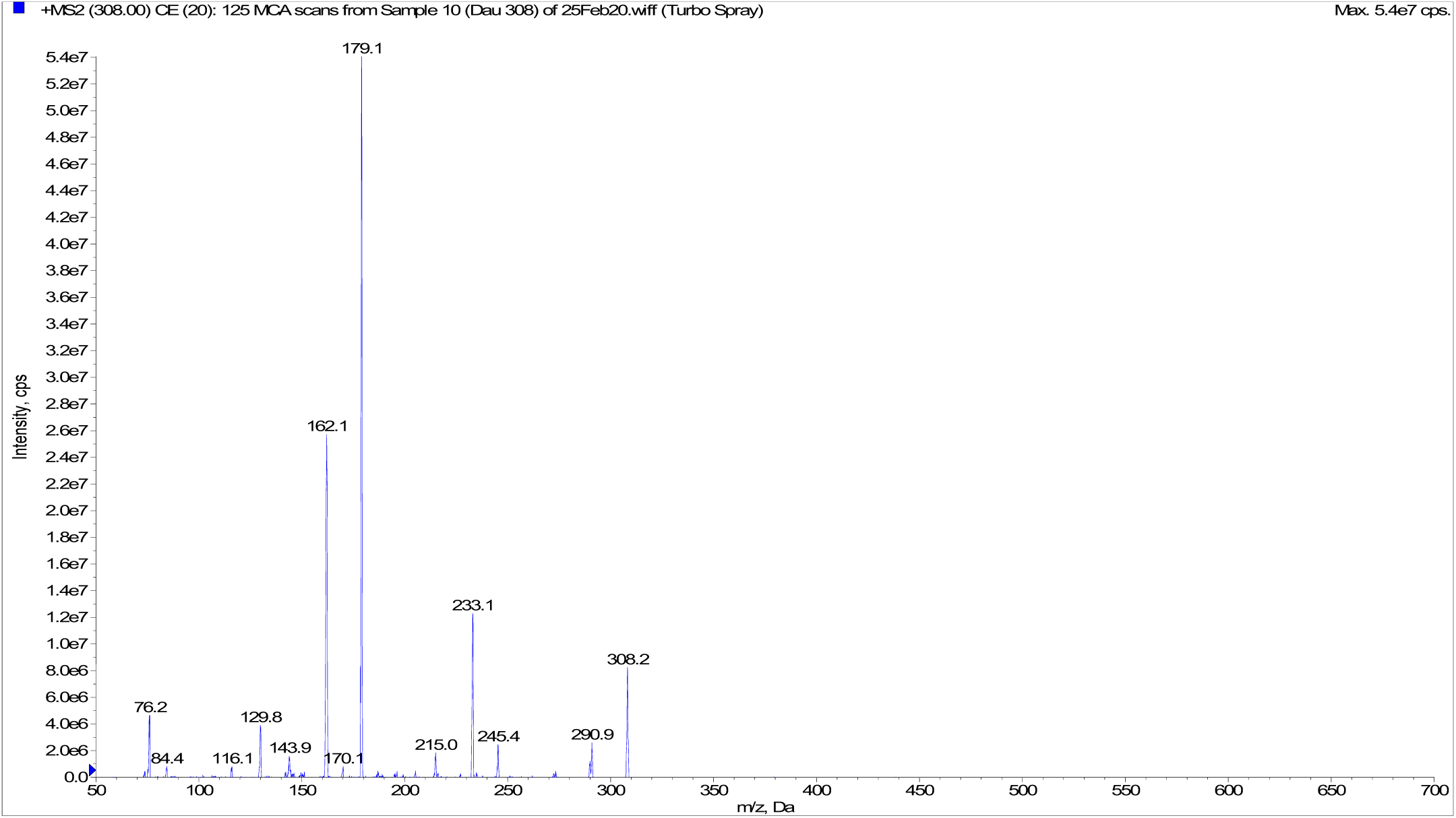
MS/MS spectra for GSH

**Figure 7.**
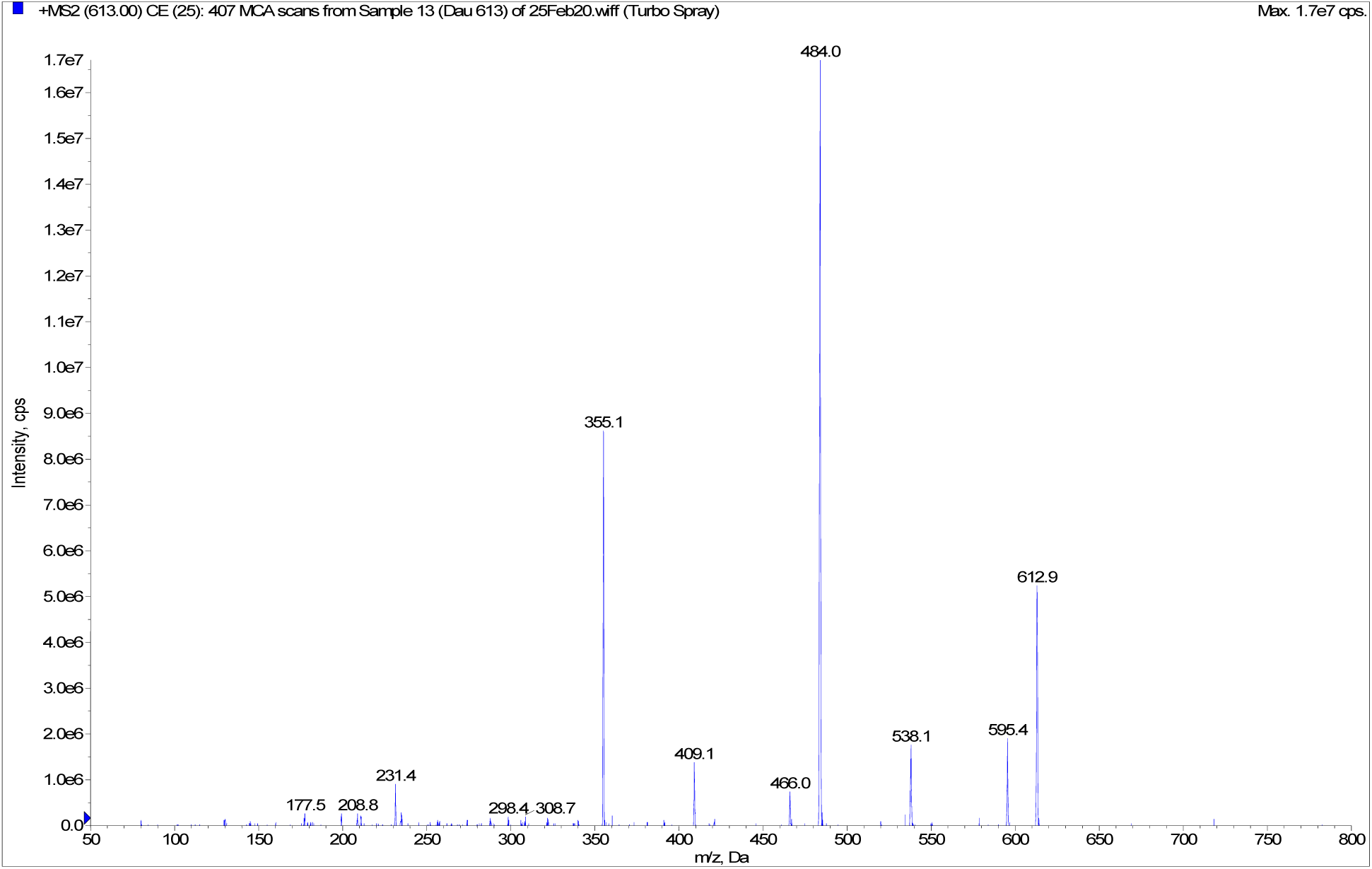
MS/MS spectra for GSSG

We made chromatograms using ion intensity (Figures 8, 9).

**Figure 8.**
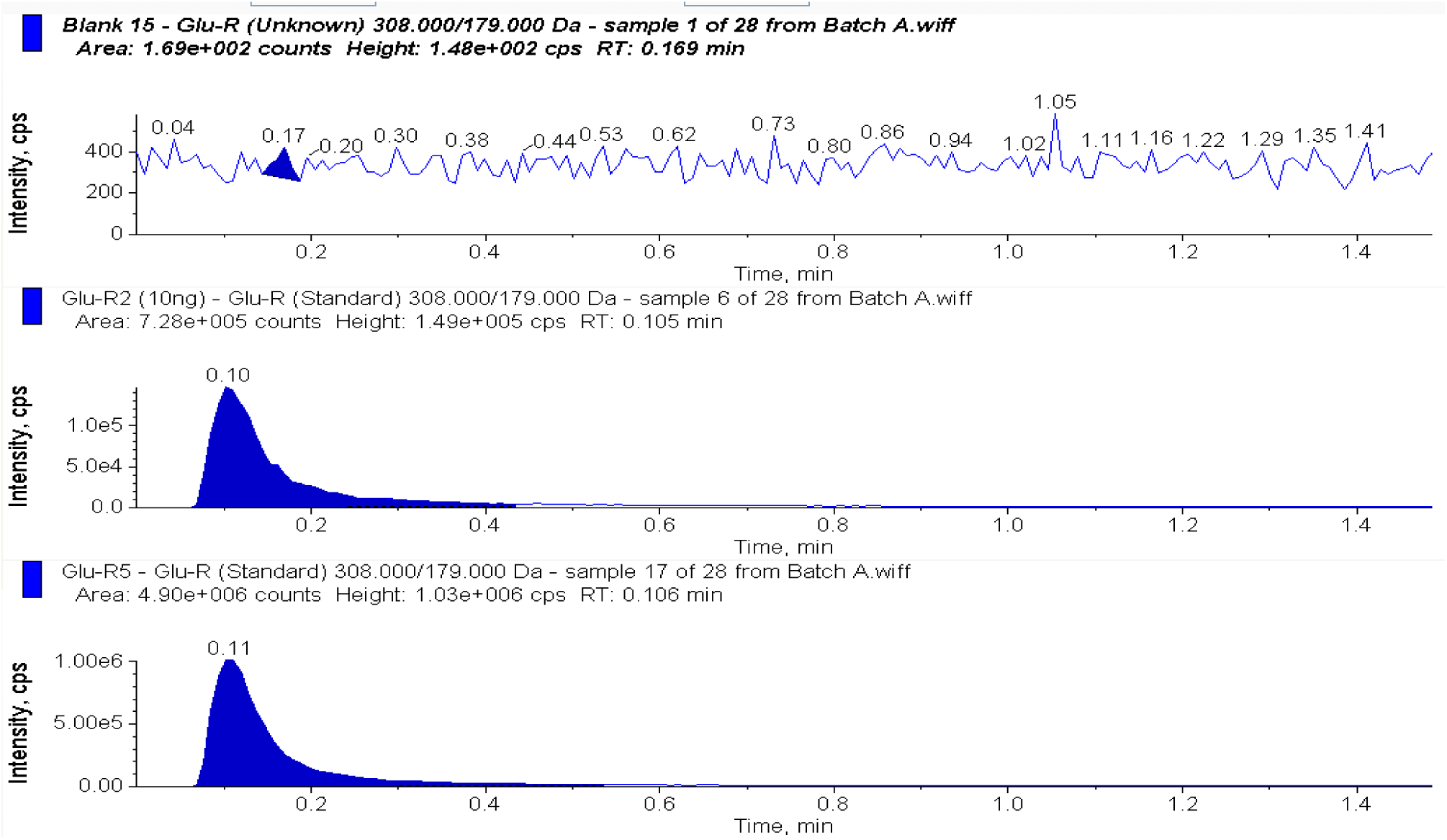
Representative chromatograms of (A) blank, (B) 10ng GSH and (C) 1000 ng GSH calibration standard

**Figure 9.**
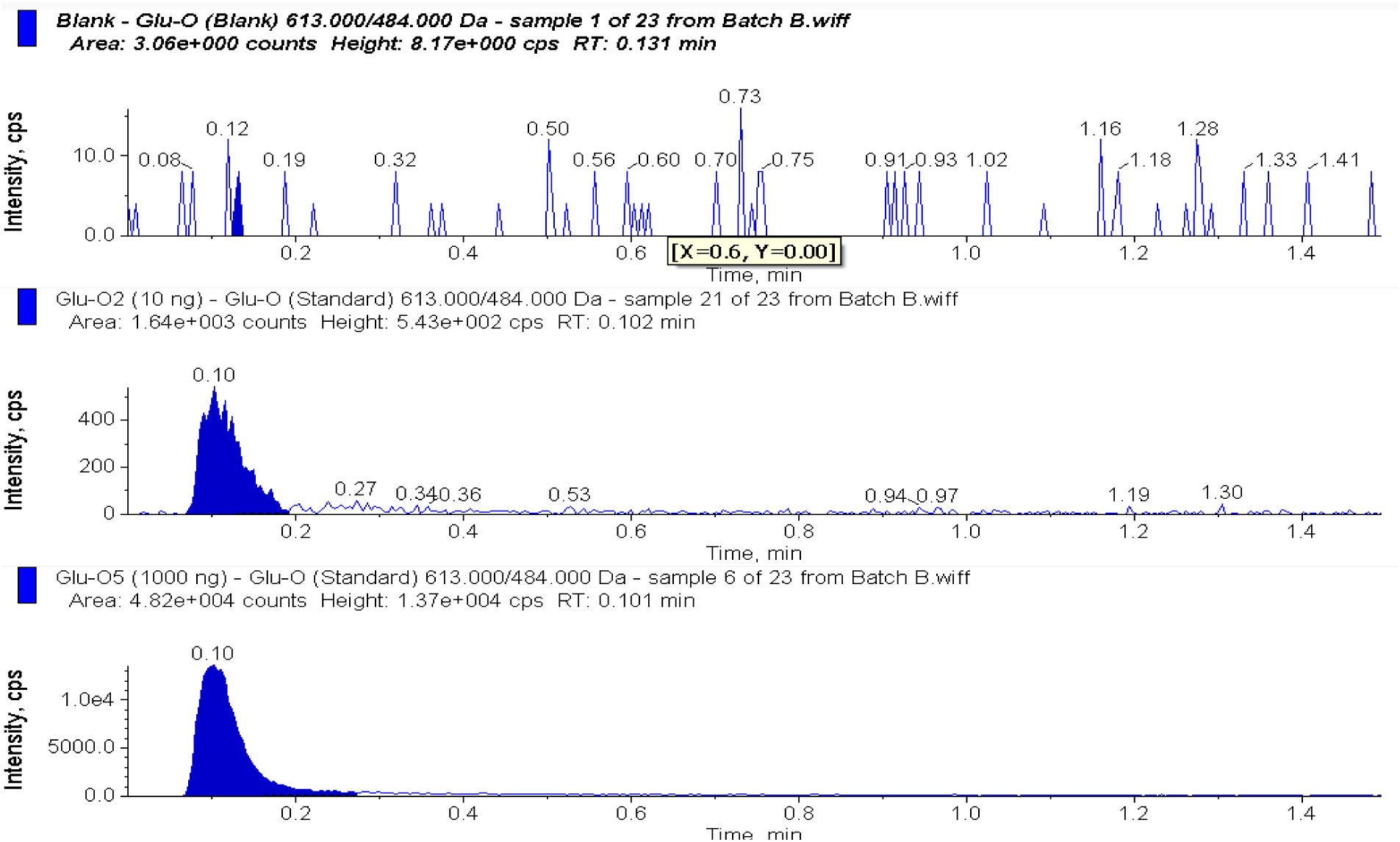
Representative chromatograms of (A) blank, (B) 10ng GSSG and (C) 1000 ng GSSG calibration standard

We used the peak areas in the chromatograms to make calibration curves. Figure 10 shows a representative calibration cure for GSSG.

**Figure 10.**
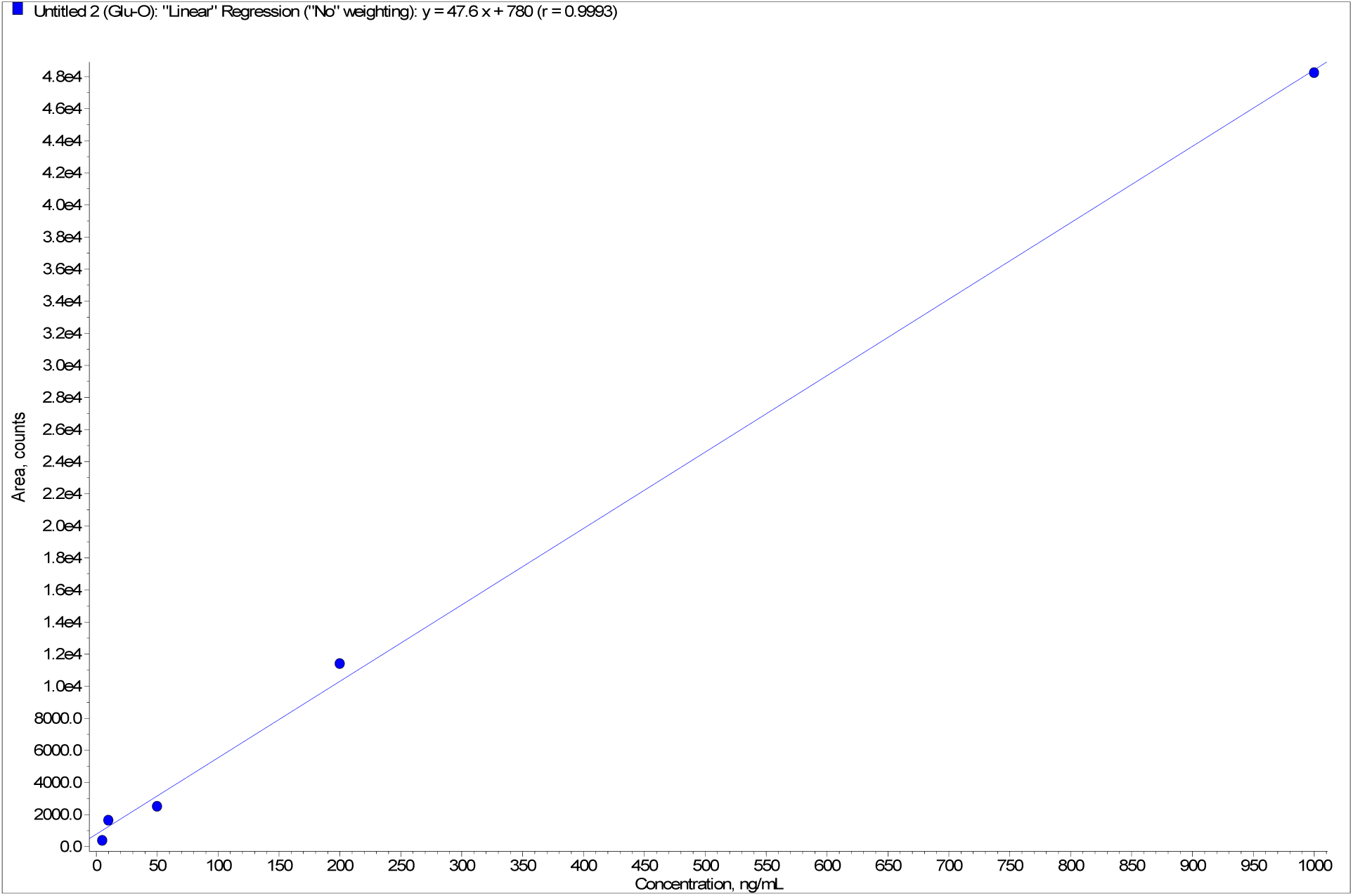
Representative of the standard calibration curve for GSSG

## Conclusion

Although there are several methods available to test for the GSH to GSSG ratio, all have disadvantages including the need to generate derivatives, the inability to conveniently measure both GSH and GSSG, and lack of enough sensitivity to allow detection in very small samples. This method solves all these problems and creates an extremely fast and reliable method. However, in our original method, and first test, the glutathione standard was not stable. We overcame this challenge by adding formic acid to the solution and storing it at 4°C as soon as we were done preparing the sample, giving us stability for at least 2 weeks. A quantitative procedure for the determination of GSH and GSSG, over the concentration range of 5.00–1,000 ng/mL, has been successfully developed and proved to be sensitive, specific, linear, precise and accurate. The method utilized a sample volume of 0.0100 mL. We developed a HPLC-MS/MS method that is fast, selective, highly sensitive, and multiple compound detectable that has the potential to be used as potential biomarker for cancer.

## Future work

We will perform flow cytometry for single cell analysis to develop a simple approach to correlate the immune cell and glutathione. The correlation of immune cells and glutathione level in blood can be used as a screening tool. There have been no studies exploring it, therefore, we purpose to develop an assay to address this correlation with potential to be used as biomarker model for cancer screening.

